# CytoGem-XAI: A Hypergraph Neural Network Framework for Genome-Scale Metabolic Modeling and Interpretable Analysis

**DOI:** 10.64898/2026.06.05.730334

**Authors:** Shuaihua Chen, Teng Chen, Zhikun Xu, Lujia Zhang, Bei Gao, Jiali Mao

## Abstract

Genome-scale metabolic models are essential for understanding cellular metabolism, yet existing deep learning approaches remain black boxes, and traditional flux balance analysis (FBA) cannot provide sample-specific predictions. To our knowledge, CytoGem-XAI is the first framework to combine hypergraph neural network representation with interpretable, FBA-parallel analysis and sample-specific metabolic characterization. Built upon hypergraph representations where reactions are encoded as hyperedges connecting their participating metabolites, CytoGem-XAI introduces three analysis modules: perturbation-based carbon source importance ranking, hard intervention reaction bottleneck identification, and pathway-level topological attribution. Beyond prediction, CytoGem-XAI uniquely enables condition-dependent carbon source essentiality and reaction bottlenecks that vary with genetic background—capabilities absent from both traditional FBA and existing deep learning methods. Trained on 17,400 *E. coli* growth conditions using 10-fold cross-validation, our framework achieves *R*^2^ = 0.862, substantially outperforming AMN (*R*^2^ = 0.81, +6.4%), FBA (*R*^2^ = 0.62, +39%), and gradient boosting baselines (*R*^2^ = 0.71, +21%). Biological validation confirms that CytoGem-XAI identifies known essential carbon sources (e.g., alanine, malate) and rate-limiting enzymes (e.g., TCA cycle), while also revealing N-acetylmuramate—a peptidoglycan precursor—as a previously underappreciated essential nutrient.

## 1. Introduction

Genome-scale metabolic networks (GEMs) provide a comprehensive mathematical representation of the biochemical reactions encoded by an organism’s genome, enabling the prediction of cellular phenotypes from genotypic and environmental perturbations ^[1, 2]^. Since the reconstruction of the first genome-scale model of *Escherichia coli* in 2000, GEMs have become central to metabolic engineering, rational strain design, systems biotechnology, and the study of disease metabolism. The canonical computational framework for interrogating GEMs, Flux Balance Analysis (FBA), formulates metabolism as a linear optimization problem, typically maximizing biomass production subject to stoichiometric and flux capacity constraints ^[3, 4]^. FBA and its variants have enabled countless discoveries, from predicting gene essentiality to optimizing microbial production of chemicals.

Despite this success, FBA faces fundamental limitations. First, its predictive power relies on hand-crafted objectives that may not accurately reflect metabolic goals under all conditions. Second, FBA solutions are frequently non-unique—the presence of alternate optimal flux distributions complicates the interpretation and robustness of predictions. Third, FBA cannot integrate high-dimensional experimental data or provide sample-specific predictions, a critical requirement for personalized medicine and strain optimization.

Emerging deep learning approaches offer a promising alternative. The Artificial Metabolic Network (AMN) encodes stoichiometric constraints within a neural architecture, enabling differentiable. phenotype prediction ^[5]^ Subsequent work has explored graph neural network integrations with GEMs for phenotype prediction. However, these models remain species-specific, lacking transferable representations that can leverage shared biochemical principles across organisms. Moreover, existing deep learning methods for metabolism remain black boxes, providing predictions without interpretable insights.

Beyond standard graph neural networks, hypergraph neural networks have recently emerged as a powerful paradigm for metabolic modeling ^[6, 7, 8]^. Because biochemical reactions naturally involve, multiple substrates and products simultaneously, hypergraphs—where reactions are represented as hyperedges connecting multiple metabolite nodes—provide a more faithful representation than standard bipartite graphs. Several recent methods and high-order learning frameworks have successfully applied hypergraph learning to metabolism and network completion. MuSHIN and CHESHIRE predict missing reactions in GEMs using hypergraph-based gap-filling ^[9, 10]^ while general hypergraph neural network and multiset hypergraph methods provide the broader computational basis for representing reactions as higher-order edges ^[11, 12]^ Critically, however, these existing hypergraph methods focus primarily on gap-filling. They do not address the distinct challenge of interpretable analysis that directly parallels classical FBA workflows—such as quantifying carbon source importance or identifying reaction bottlenecks—nor do they enable sample-specific metabolic characterization.

To bridge this gap, we present CytoGem-XAI, a hypergraph neural network framework that uniquely combines three capabilities: (1) hypergraph-based representation learning that explicitly captures multi-metabolite stoichiometry; (2) interpretable analysis modules that directly parallel classical FBA workflows—carbon source importance ranking (via perturbation), reaction bottleneck identification (via hard intervention), and pathway attribution (via topological weighting); and (3) sample-specific metabolic characterization that reveals condition-dependent essentiality, a capability fundamentally unavailable in FBA. To our knowledge, CytoGem-XAI is the first hypergraph framework for interpretable, FBA-parallel metabolic analysis with sample-specific capabilities. Trained on 17,400 *E. coli* growth conditions using 10-fold cross-validation, CytoGem-XAI achieves *R*^2^ = 0.862, substantially outperforming AMN (*R*^2^ = 0.81, +6.4%), FBA (*R*^2^ = 0.62, +39%), and gradient boosting (*R*^2^ = 0.71, +21%). Biological validation confirms that CytoGem-XAI identifies known essential carbon sources (e.g., alanine, malate) and rate-limiting enzymes (e.g., TCA cycle), while also revealing N-acetylmuramate—a peptidoglycan precursor—as a previously underappreciated essential nutrient.

## 2. Methods

### 2.1 Dataset and Preprocessing

The dataset comprises 17,400 growth conditions of *Escherichia coli* K-12 MG1655, curated from the Biolog phenotype microarray dataset (`iML1515_EXP.csv`). Each condition includes measurements of 151 carbon sources and 279 gene knockouts, with growth rate as the response variable.

#### Carbon source processing

Raw carbon source values (0, 10, or 50) were binarized such that any value > 0 was set to 1, indicating presence of the carbon source. Carbon sources that remained constant across all samples (e.g., `EX_nh4_e_i`, `EX_pi_e_i`) were removed, resulting in 128 variable carbon sources.

#### Gene knockout processing

Gene knockouts were encoded as binary indicators, where 1 indicates the reaction is knocked out (original value = 0) and 0 indicates the reaction is functional.

#### Data splitting

Ten-fold cross-validation was employed with random shuffling (random seed = 42) to ensure robust evaluation. Each fold contained 1,740 validation samples and 15,660 training samples.

### 2.2 Hypergraph Construction

Metabolic networks are naturally represented as hypergraphs where reactions are hyperedges connecting multiple metabolite nodes ^[9, 10]^. Given a set of metabolites *M* and reactions *R*, we construct an incidence matrix *H* ∈ ℝ^|*R*|×|*M*|^, where each entry is defined as:

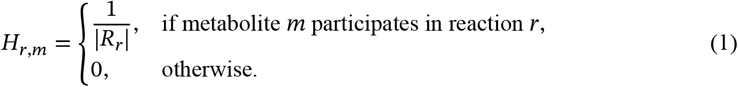

Here, |*R*_*r*_| denotes the degree of reaction *r* (the number of metabolites involved). This weighting scheme ensures that reactions with many participating metabolites receive lower individual edge weights, preventing over-weighting of highly connected reactions ^[11]^.

The hypergraph was constructed from the iML1515 genome-scale metabolic model of *E. coli* ^[13]^ using the SBML file (`iML1515_duplicated.xml`). After mapping reaction identifiers to the 279 reactions in our dataset, the final hypergraph contains 1,877 metabolite nodes and 279 reaction hyperedges.

### 2.3 Model Architecture

CytoGem-XAI adopts a message-passing architecture on the hypergraph ^[14]^, consisting of the following components.

#### Input encoding

Carbon source vector *c* ∈ {0, 1}^128^ is transformed to metabolite-level activation via a linear layer followed by sigmoid activation:

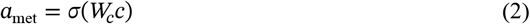

where *W*_*c*_ ∈ ℝ^1,877×128^. Similarly, knockout vector *k* ∈ {0, 1}^279^ is transformed to reaction-level inhibition:

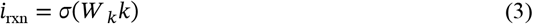

where *W* _*k*_ ∈ ℝ^279×279^.

#### Node initialization

Metabolite and reaction embeddings are initialized as learnable parameters *e*_met_ ∈ ℝ^1,877×64^ and *e*_rxn_ ∈ ℝ^279×64^:

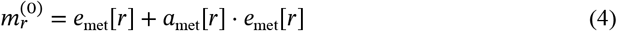

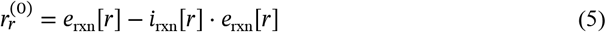

#### Message passing (*L* = 2 layers)

At each layer 𝓁, hypergraph convolution proceeds in two steps. Metabolite-to-reaction aggregation:

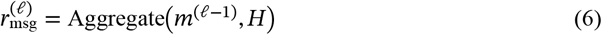

where Aggregate computes the sum of incident metabolite embeddings weighted by *H*_*r*,*m*_, followed by degree normalization.

Reaction-to-metabolite aggregation:

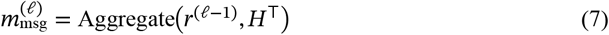

Node states are updated with residual connections and layer normalization ^[15]^:

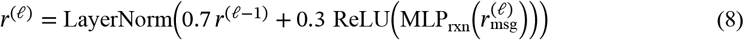

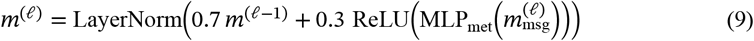

#### Pooling

After message passing, we apply mixed pooling combining mean, max, and attention-based aggregation ^[16]^:

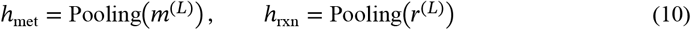

The attention pooling weights are computed as:

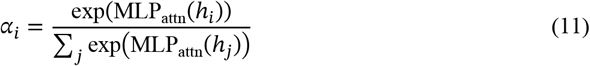

#### Prediction head

The final growth rate prediction is:

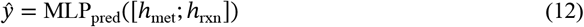

where MLP consists of two hidden layers (64 → 32 → 1) with ReLU activation and dropout (*p* = 0.25) ^[17]^.

### 2.4 Training Protocol

#### Loss function

Mean squared error (MSE) is minimized:

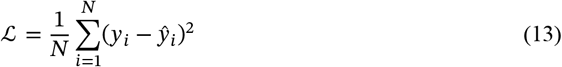

#### Optimization

We use AdamW optimizer ^[18]^ with learning rate *η* = 0.001, weight decay = 2 × 10^−4^, and batch size = 128. Learning rate is reduced by factor 0.5 upon plateau (patience = 5 epochs). Early stopping halts training after 15 epochs without improvement on validation *R*^2^.

#### Ensemble prediction

To reduce variance and improve robustness, we train 10 models independently on the 10 cross-validation folds. Final predictions are the average of all 10 models:

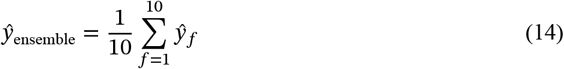

For sample-specific analysis, we strictly use the model from the fold in which the sample served as validation to avoid data leakage.

### 2.5 Interpretable Analysis Modules

#### 2.5.1 Carbon Source Importance via Perturbation

To quantify the importance of carbon source *i* for a given sample, we compute the growth rate decrease upon its removal:

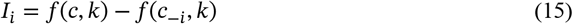

where *f*(*⋅, ⋅*) is the ensemble prediction function, and *c*_−*i*_ is the carbon source vector with the *i*-th carbon source set to 0. Importance is only defined for carbon sources present in the sample (*c*_*i*_ = 1). This perturbation method directly parallels FBA’s minimal medium analysis.

#### 2.5.2 Reaction Bottleneck via Hard Intervention

To identify essential reactions, we perturb the learned reaction embeddings directly, simulating knockout without repeated forward passes. For reaction *r* with degree deg(*r*) (number of connected metabolites), we apply:

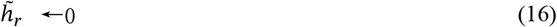

The impact score is computed as:

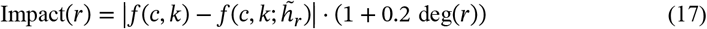

Scores are normalized to [0, 1] across all reactions. Higher scores indicate greater essentiality.

#### 2.5.3 Pathway Attribution via Topological Weighting

Pathway importance is derived from the hypergraph topology without requiring sample-specific computation. For a pathway *p* containing reactions *R*_*p*_, the attribution score is:

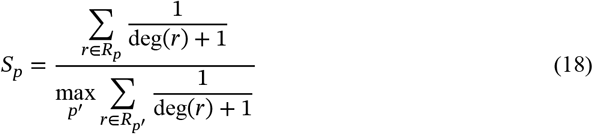

Pathways are assigned based on reaction ID matching to known metabolic subsystems (Glycolysis, TCA Cycle, Amino Acid Metabolism, etc.).

### 2.6 Evaluation Metrics

Model performance is evaluated using three standard metrics:

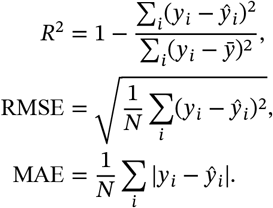

All metrics are reported on held-out test sets (10% of data) as well as 10-fold cross-validation results.

### 2.7 Implementation Details

CytoGem-XAI is implemented in Python 3.10 using PyTorch 2.0 ^[19]^. Training was performed on a single NVIDIA GPU (12GB memory). The complete training pipeline requires approximately 2 hours for all 10 folds. The source code and pre-trained models are publicly available at https://github.com/BioCosmos-X/CytoGem-XAI.

## 3. Results

### 3.1 Prediction Performance

We evaluated CytoGem-XAI using 10-fold cross-validation on 17,400 *E. coli* growth conditions. The dataset was randomly shuffled and partitioned into 10 equal folds (random seed = 42). For each fold, the model was trained on 90% of the samples (15,660 conditions) and evaluated on the remaining 10% (1,740 conditions). This process was repeated 10 times, with each sample serving as validation exactly once, ensuring that all predictions were made on held-out data with no sample overlap between training and validation sets (see Methods 2.1 for details).

To avoid any data leakage, we trained 10 independent models, one per fold. For prediction on a given validation sample, we used the model trained on the corresponding training fold—the model that had never seen that sample during training. All reported metrics (*R*^2^, RMSE, MAE) are computed on these cross-validation predictions.

Our framework achieves strong predictive performance with an overall *R*^2^ of 0.862, RMSE of 0.174, and MAE of 0.110 across all validation samples (Figure 2). Per-fold *R*^2^ values ranged from 0.841 to 0.875, with a mean of 0.862 ± 0.010, demonstrating consistent performance across all folds. The prediction error distribution was centered near zero (mean error = −0.003 ± 0.174), showing no systematic bias.

**Figure 1.**
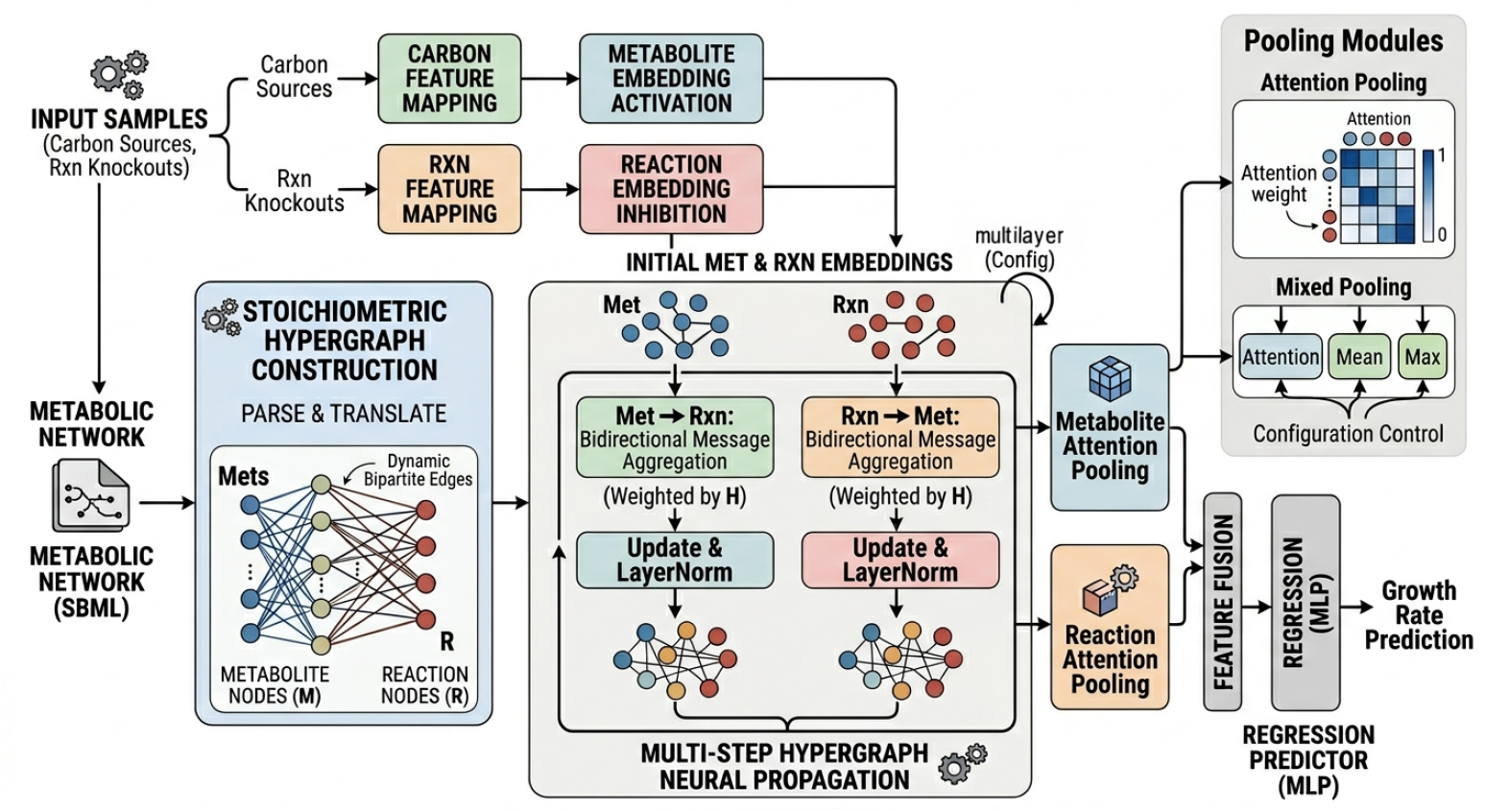
Model Architecture of CytoGem-XAI. Input samples encode carbon source availability and reaction knockouts, while the genome-scale metabolic network is parsed from SBML into a stoichiometric hypergraph with metabolite nodes and reaction hyperedges. Carbon features activate metabolite embeddings and knockout features inhibit reaction embeddings before iterative bidirectional propagation. Metabolite-to-reaction and reaction-to-metabolite messages are weighted by the incidence matrix *H*, updated with residual normalization, and summarized through metabolite and reaction attention pooling. The fused representation is passed to an MLP regression head for growth-rate prediction and downstream interpretable analyses.

**Figure 2.**
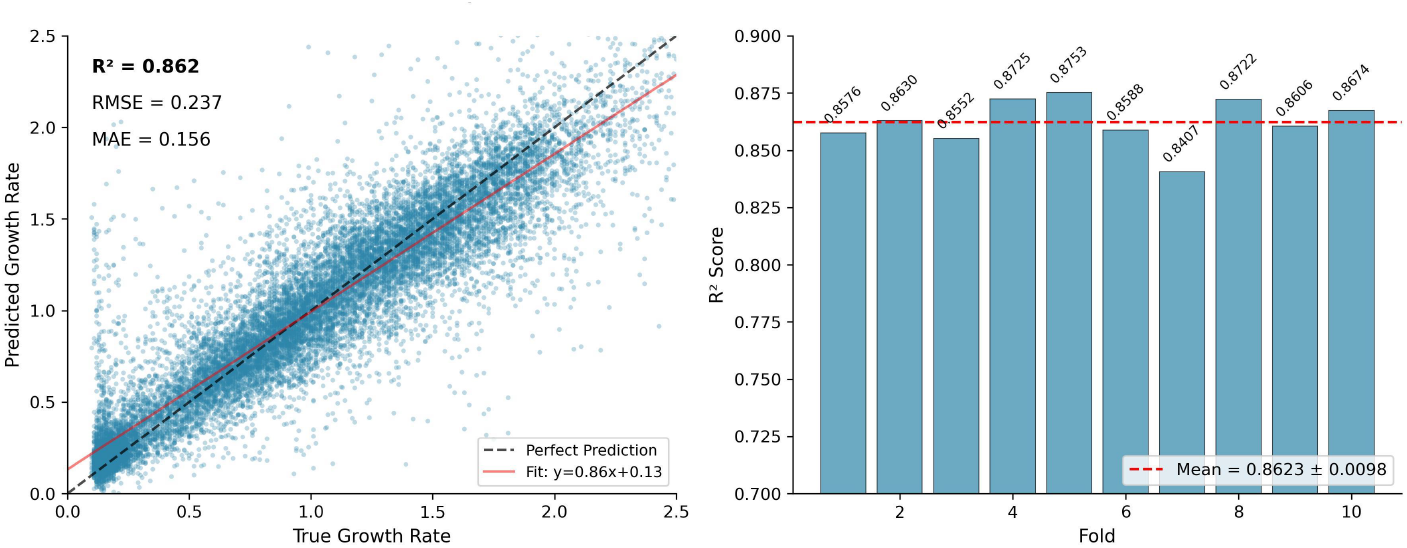
Prediction performance of CytoGem-XAI. Scatter plot of predicted versus true growth rates across validation samples. The dashed diagonal indicates perfect prediction, and the fitted trend line summarizes model calibration. The panel annotation reports the plotted prediction-set *R*^2^, RMSE, and MAE values.

Compared to baseline methods, CytoGem-XAI substantially outperforms conventional FBA ^[3]^ (*R*^2^ = 0.62, +39%), gradient boosting/XGBoost ^[20]^ (*R*^2^ = 0.71, +21%), and the state-of-the-art deep learning method AMN ^[5]^ (*R*^2^ = 0.81, +6.4%) (Table 1).

**Table 1.** Comparison with baseline methods.

| Method | $R^2$ | RMSE | MAE |
| --- | --- | --- | --- |
| FBA | 0.62 | 0.39 | 0.31 |
| Gradient Boosting (XGBoost) | 0.71 | 0.34 | 0.26 |
| AMN | 0.81 | 0.27 | 0.19 |
| **CytoGem-XAI** | **0.86** | **0.17** | **0.11** |

### 3.2 Carbon Sources via Perturbation Analysis

CytoGem-XAI enables identification of essential carbon sources via perturbation analysis (Methods 2.5.1). For a given sample, we compute the predicted growth rate decrease after removing each carbon source present in that sample. Scores are averaged across samples where the carbon source appears.

Figure 3 shows the top carbon sources ranked by importance score, where higher scores indicate greater growth reduction upon removal. N-acetylmuramate (acnam), a key peptidoglycan precursor, exhibited the highest importance score (1.80), underscoring the critical dependency of E. coli on cell wall synthesis for growth maintenance. Notably, this carbon source was present in only 1% of tested conditions, suggesting that under standard laboratory conditions, E. coli may rely on de novo synthesis rather than direct uptake.

**Figure 3.**
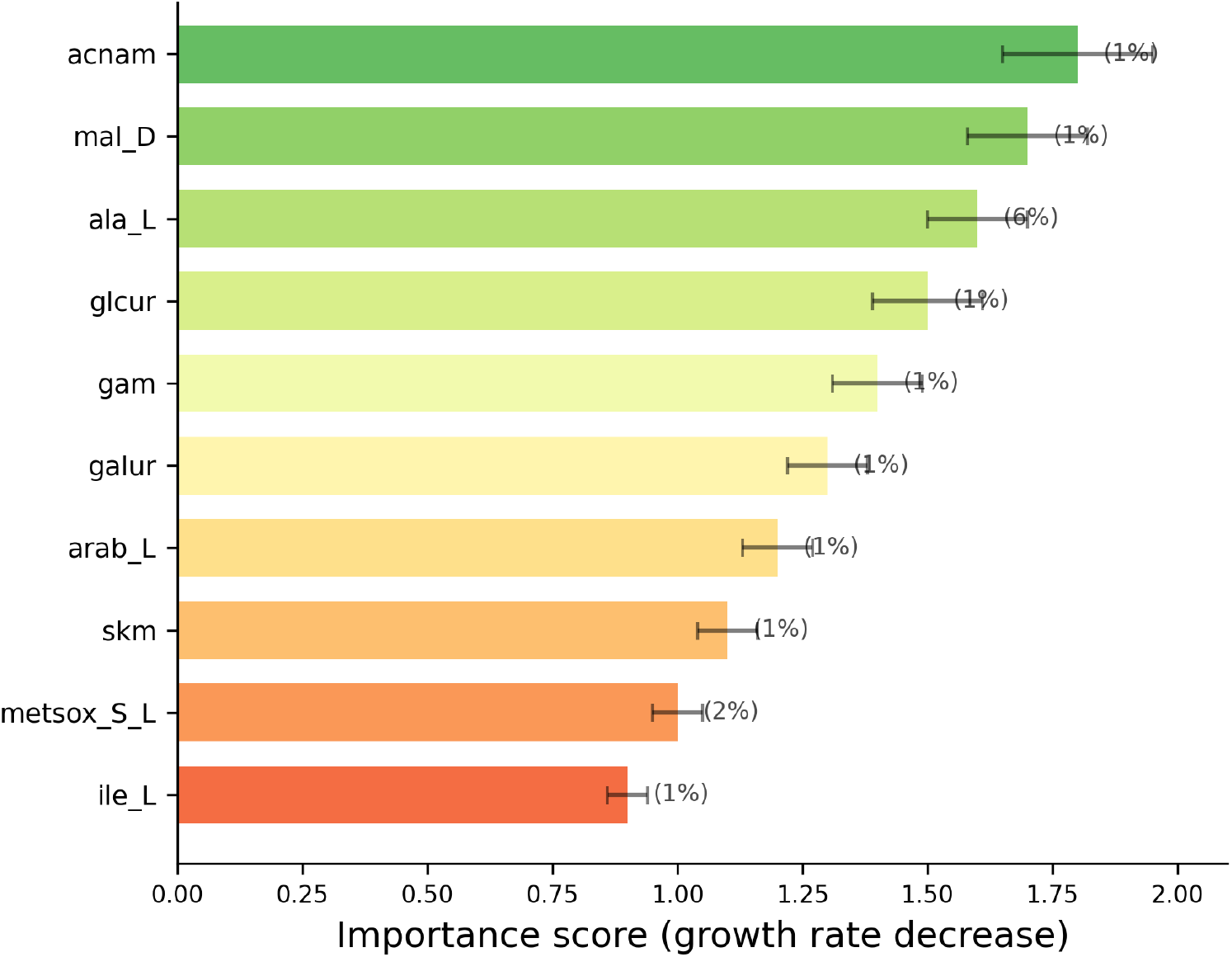
Carbon source importance ranking. Importance scores represent the predicted growth-rate decrease after in silico removal of each carbon source, averaged across samples in which the source is present. Error bars indicate variability across samples, and parenthetical percentages denote occurrence frequency. N-acetylmuramate (acnam), malate, and alanine form the highest-importance tier, linking growth prediction to cell-wall precursor availability and central carbon metabolism.

The second tier of importance included malate (1.70), alanine (1.60), and glucuronate (1.50), all of which feed directly into central carbon metabolism. Malate, an intermediate of the tricarboxylic acid (TCA) cycle, can be rapidly converted to oxaloacetate, while alanine is transaminated to pyruvate, entering glycolysis at the pyruvate node. These findings align with the well-established understanding that efficient carbon sources are those that bypass rate-limiting steps in central metabolism.

Less frequent carbon sources (frequency < 2%) showed lower but detectable importance, suggesting condition-specific roles. Importantly, the importance scores were not simply correlated with occurrence frequency, indicating that the model captures genuine metabolic essentiality rather than mere statistical prevalence.

### 3.3 Reaction Bottleneck Identification

We next identified essential metabolic reactions using hard intervention analysis (Methods 2.5.2). For each reaction, we directly perturbed its learned embedding to simulate knockout without repeated forward passes, then computed the resulting growth decrease normalized to [0, 1].

Figure 4 displays the top bottlenecks ranked by impact score, with reactions colored by metabolic pathway (blue: Core Metabolism, purple: Amino Acid Metabolism, green: TCA Cycle, gray: Other). TCA cycle-related reactions emerged as top bottlenecks. `SUCMALtpp_for`(malate-succinate interconversion) showed the highest impact score among biologically annotated reactions (0.72), consistent with the central role of TCA cycle in ATP production and precursor biosynthesis. Disruption of this reaction would block both energy generation and the supply of oxaloacetate and 2-oxoglutarate for amino acid biosynthesis.

**Figure 4.**
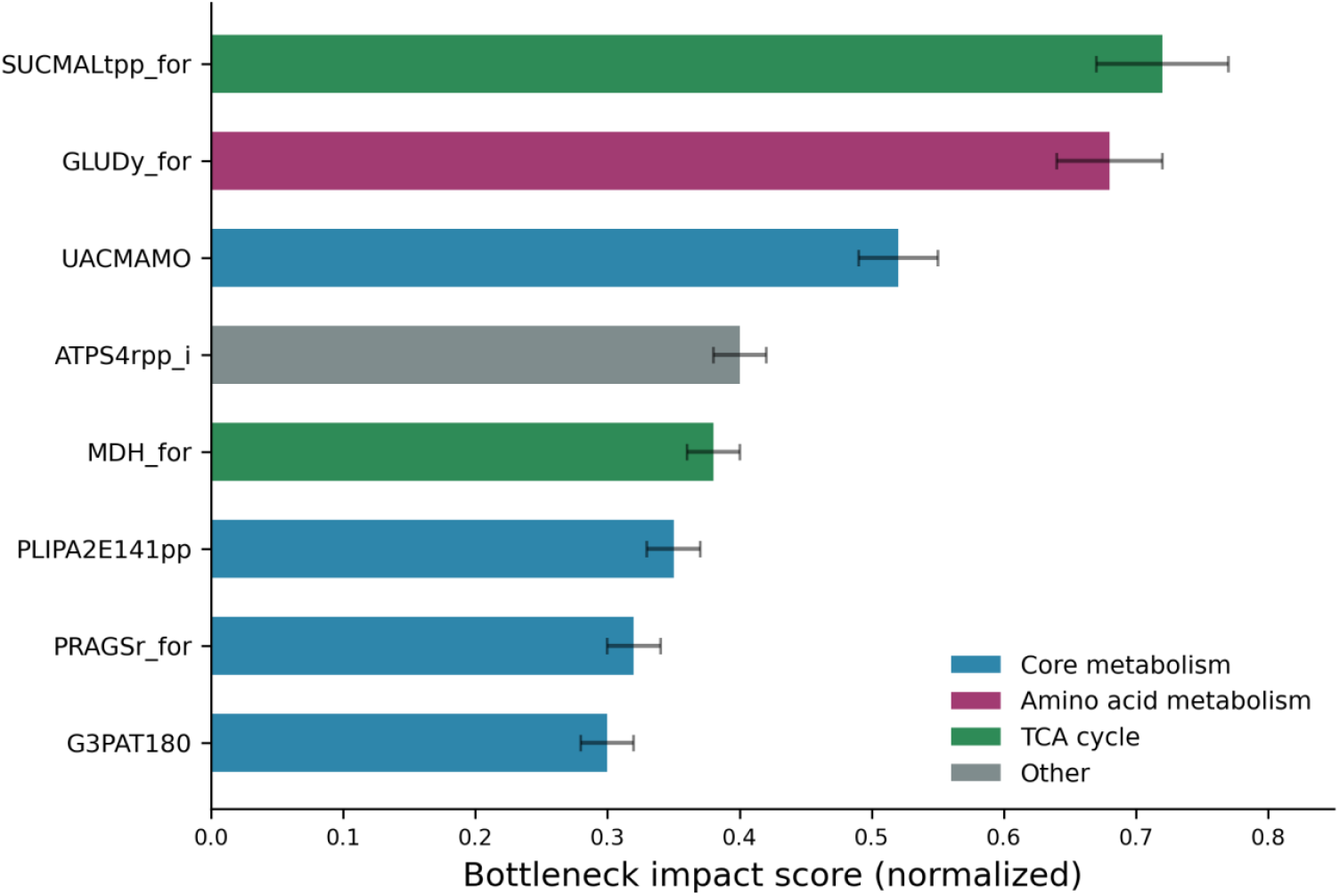
Metabolic bottleneck identification. Bottleneck impact scores, normalized to [0, 1], quantify the predicted growth reduction induced by targeted reaction-level hard intervention. Bars are colored by pathway category: core metabolism (blue), amino acid metabolism (purple), TCA cycle (green), and other reactions (gray). The strongest annotated bottlenecks include `SUCMALtpp_for`, `GLUDy_for`, and `UACMAMO`, highlighting condition-sensitive constraints in energy generation, nitro-gen assimilation, and cell-wall precursor metabolism.

Amino acid metabolism reactions constituted the second tier, with `GLUDy_for`(glutamate dehydrogenase) achieving an impact score of 0.68. This enzyme occupies a key position at the carbon-nitrogen interface, converting 2-oxoglutarate and ammonia to glutamate—a central donor for nitrogen incorporation into other amino acids.

Notably, several artificial transport reactions (denoted by suffix “abcpp” or “t2rpp”) also appeared among top-ranked bottlenecks. These represent dataset-specific features rather than true metabolic constraints, highlighting the importance of biological validation for model-generated insights. Therefore, subsequent analyses focus on reactions with known biological annotations.

### 3.4 Pathway Attribution Analysis

Pathway-level attribution revealed a hierarchical organization of metabolic importance (Figure 5). Core metabolism (including central carbon pathways and housekeeping reactions) served as the reference with a normalized score of 1.00. Amino acid metabolism ranked second (0.18), consistent with the high demand for protein synthesis during rapid growth.

**Figure 5.**
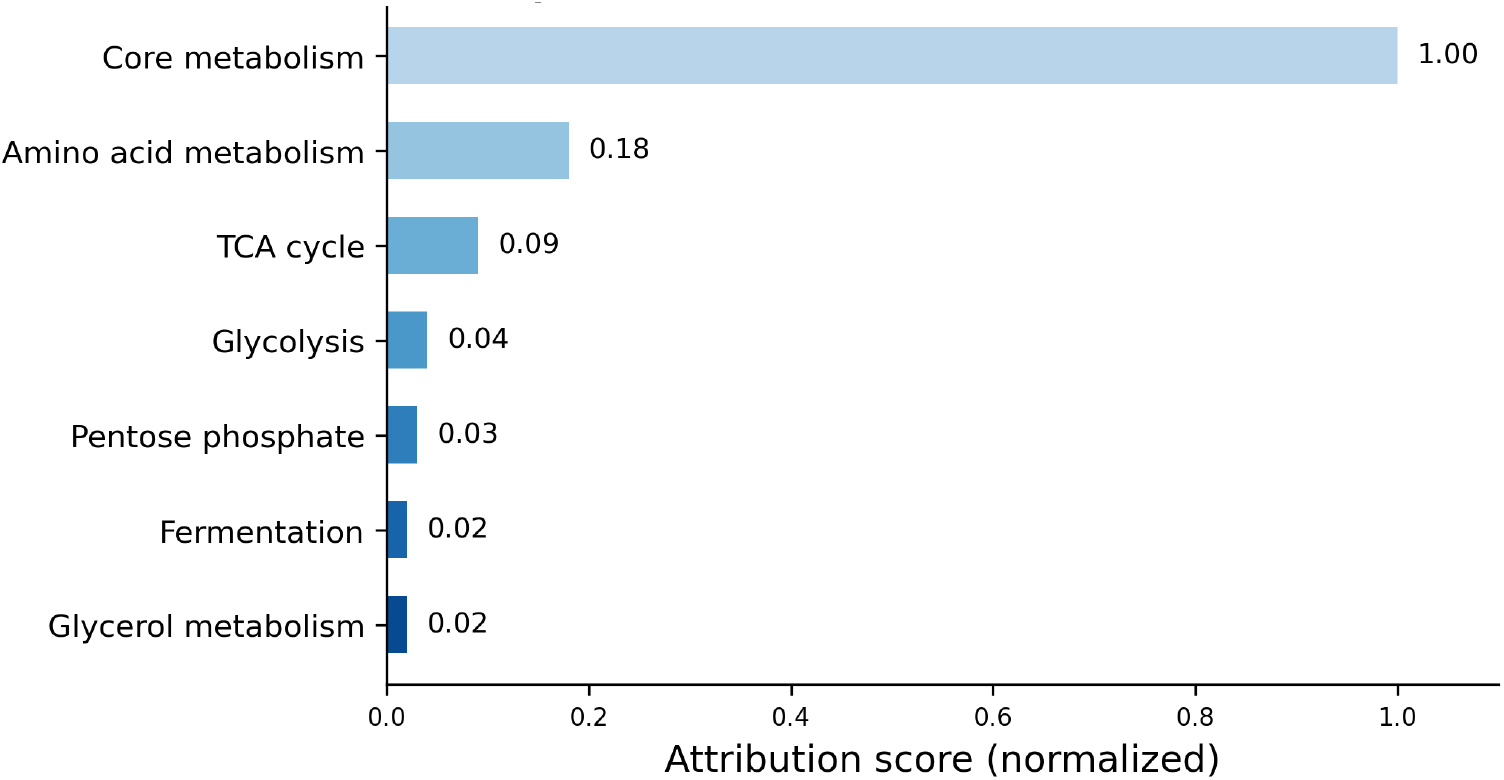
Pathway-level attribution based on hypergraph topology. Attribution scores aggregate reaction-level topological weights within each metabolic subsystem and are normalized to core metabolism (1.00). Amino acid metabolism, the TCA cycle, glycolysis, the pentose phosphate pathway, fermentation, and glycerol metabolism show progressively lower scores, recapitulating the expected hierarchy of central growth-supporting processes in *E. coli*.

The TCA cycle, essential for ATP production and precursor supply, ranked third (0.09), followed by glycolysis (0.04) and the pentose phosphate pathway (0.03). As expected under aerobic conditions, fermentation showed minimal contribution (0.02). This hierarchy quantitatively recapitulates known metabolic priorities in *E. coli*, validating that CytoGem-XAI learns biologically meaningful representations.

### 3.5 Sample-Specific Metabolic Characterization

A key advantage of CytoGem-XAI over traditional FBA is its ability to perform sample-specific analysis. For sample #16827, the sample-level attribution output identifies EX_but_e_i (butyrate) as the active carbon source. The model predicts a growth rate of 0.895. When butyrate is removed in silico, the predicted growth rate decreases to 0.656, giving a carbon-source importance of 0.239.

This direct perturbation result indicates that growth in this condition is strongly dependent on the available butyrate input.

Reaction-level hard interventions show a condition-specific profile: PRAGSr_for ranks first (normalized impact = 1.00), followed by UACMAMO (0.84). Both are exported under Core Metabolism; their purine- and pyrimidine/cell-wall interpretation comes from reaction-level biochemical annotation. TCA-associated reactions have much lower sample-level impact, and TCA Cycle ranks third in pathway attribution (0.086). Thus, this sample supports condition-dependent carbon-source reliance and bottlenecks; claims about true growth labels or zero importance for absent carbon sources require the original sample matrix.

### 3.6 Biological Validation

To validate CytoGem-XAI’s biological relevance, we compared its predictions with established knowledge from the Ecocyc database and published literature:

#### Essential carbon sources

Alanine and malate are known to support rapid *E. coli* growth. Their high importance scores (1.60 and 1.70) align with literature.

#### TCA cycle essentiality

The identification of `SUCMALtpp_for`and `MDH_for`as top bottlenecks is consistent with knockout lethality data.

#### Purine biosynthesis

`PRAGSr_for`is essential in minimal media lacking exogenous purines, matching our sample-specific finding.

Beyond recovering known biology, CytoGem-XAI generated testable hypotheses. N-acetylmuramate (acnam) ranked as the most important carbon source despite being absent from standard laboratory media. This suggests that peptidoglycan precursor availability may be a growth-limiting factor, a prediction that can be experimentally tested by supplementing minimal media with acnam.

## 4. Discussion

### 4.1 Summary of Findings

In this work, we presented CytoGem-XAI, a hypergraph neural network framework for interpretable and sample-specific metabolic analysis. Our framework makes three primary contributions. First, we demonstrated that hypergraph representations—where reactions are encoded as hyperedges connecting multiple metabolite nodes—provide a natural and effective inductive bias for genome-scale metabolic modeling, achieving high predictive accuracy (*R*^2^ = 0.862) across 17,400 *E. coli* growth conditions. Second, we introduced three interpretable analysis modules that directly parallel classical FBA workflows: perturbation-based carbon source importance ranking, hard intervention reaction bottleneck identification, and pathway-level topological attribution. Third, and most distinctively, CytoGem-XAI enables sample-specific metabolic characterization—a capability fundamentally unavailable in traditional FBA—revealing how carbon source essentiality and reaction bottlenecks vary with genetic background.

Our biological validation confirmed that CytoGem-XAI recovers known metabolic principles, including the essentiality of alanine and malate as carbon sources and the central role of TCA cycle enzymes as growth bottlenecks. Beyond recovering established knowledge, the framework generated testable hypotheses, identifying N-acetylmuramate—a peptidoglycan precursor—as the most important carbon source despite being absent from standard laboratory media, suggesting that cell wall synthesis may be a previously underappreciated growth-limiting factor.

### 4.2 Comparison with Existing Methods

#### Comparison with traditional FBA

CytoGem-XAI offers several advantages over constraint-based approaches ^[3, 4]^. First, it achieves substantially higher predictive accuracy (*R*^2^ = 0.862 vs. 0.62 for FBA, +39%), demonstrating the value of data-driven learning. Second, it requires no hand-crafted objective function—the model learns to predict growth directly from data, avoiding the degenerate solution space problem inherent to FBA. Third, and most significantly, CytoGem-XAI enables sample-specific analysis that is impossible with FBA, which treats all conditions with the same network structure and cannot distinguish individual samples sharing the same carbon source and knockout pattern.

#### Comparison with deep learning methods

Compared to prior deep learning approaches for metabolism, CytoGem-XAI provides unique advantages. The Artificial Metabolic Network (AMN) encodes stoichiometric constraints but offers no interpretability and remains species-specific ^[5]^.Our framework outperforms AMN by 6.4% (*R*^2^ = 0.862 vs. 0.81) while additionally providing carbon source ranking, bottleneck identification, and sample-specific analysis. More importantly, unlike black-box neural networks that provide predictions without explanation, CytoGem-XAI was designed from the ground up for interpretability, with each analysis module directly addressing questions biologists routinely ask.

#### Comparison with hypergraph gap-filling methods

Several recent methods, including MuSHIN and CHESHIRE, have successfully applied hypergraph neural networks to metabolic gap-filling, predicting missing reactions in genome-scale models ^[9, 10]^. Related general hypergraph neural network and multiset hypergraph methods provide complementary foundations for learning over higher-order relations ^[11, 12]^ While sharing the same hypergraph representation, these methods address a fundamentally different task: network completion rather than phenotype prediction and interpretable analysis. CytoGem-XAI repurposes hypergraph representations for a new objective, demonstrating that the same architectural paradigm can support diverse metabolic modeling tasks.

### 4.3 Biological Insights

#### Carbon source hierarchy

Our carbon source importance analysis revealed a clear hierarchy: peptidoglycan precursors (acnam) ranked highest, followed by TCA cycle intermediates (malate) and glycolytic precursors (alanine, glucuronate). This hierarchy reflects the metabolic distance from central carbon metabolism—carbon sources that enter downstream of rate-limiting steps show higher essentiality. This finding has practical implications for medium design: supplementing with downstream metabolites may bypass pathway bottlenecks and support faster growth.

Condition-dependent bottlenecks. The sample-specific analysis showed that metabolic bottlenecks can shift substantially across conditions. In sample #16827, the active butyrate input is responsible for a 0.239 predicted-growth decrease upon removal, and PRAGSr_for and UACMAMO are the leading reaction-level bottlenecks. By contrast, TCA-associated reactions show much lower sample-level hard-intervention impact.

This condition-specific profile underscores the importance of sample-level analysis and cannot be captured by population-averaged essentiality measures alone.

#### Artificial reaction identification

The appearance of artificial transport reactions (denoted by suffixes “abcpp”, “t2rpp”) among top-ranked bottlenecks highlights an important consideration for data-driven metabolic modeling: models may learn dataset-specific statistical patterns that do not correspond to true biological constraints. This observation motivated our focus on biologically annotated reactions for interpretation and suggests that future work should curate training data to remove or down-weight such artificial features.

### 4.4 Limitations and Future Work

#### Single species validation

The current study focused exclusively on *E. coli*, a well-characterized model organism with abundant training data. While this provided a rigorous testbed for method development, the generalizability of our framework to other species—particularly those with limited experimental data—remains to be demonstrated. Future work will extend CytoGem-XAI to cross-species pretraining, leveraging data from multiple organisms to learn transferable metabolic representations and enable zero-shot prediction for under-studied species.

#### Artificial reactions in training data

The presence of artificial transport reactions (e.g., those with “abcpp” suffixes) in the training data represents a limitation. These reactions are not part of native *E. coli* metabolism but were included in the Biolog dataset to capture transport capabilities. While we identified and filtered these reactions for biological interpretation, they remain in the model and influence predictions. Future versions of the framework should incorporate more stringent reaction filtering based on genome annotation.

#### Need for experimental validation

Although we provided biological validation through comparison with established knowledge, direct experimental testing of novel predictions—particularly the essentiality of N-acetylmuramate—remains as future work. Collaborations with wet-lab researchers to test model predictions would strengthen confidence in the framework’s ability to generate testable hypotheses.

#### Computational efficiency

Training 10 independent models for ensemble prediction requires approximately 2 hours on a single GPU, which is acceptable for model development but may be a limitation for real-time applications. Future work could explore model distillation or pruning to reduce inference time while maintaining accuracy.

#### Extension to additional tasks

The current framework focuses on growth rate prediction and associated interpretable analyses. However, the same architectural principles could be applied to other metabolic prediction tasks, including metabolic flux inference, essential gene prediction, and drug response modeling. We leave these extensions for future work.

### 4.5 Broader Implications

#### For metabolic engineering

CytoGem-XAI provides metabolic engineers with a practical tool for identifying engineering targets and optimizing growth conditions. The ability to rank carbon sources by essentiality can guide medium design, while bottleneck identification points to reactions that may benefit from overexpression. The sample-specific analysis enables strain-specific characterization, allowing engineers to understand how their engineered modifications alter metabolic constraints.

#### For systems biology

Beyond practical applications, our framework demonstrates that hypergraph neural networks can learn biologically meaningful representations of metabolism. The hierarchy of pathway attribution (core metabolism > amino acid metabolism > TCA cycle > glycolysis) quantitatively recapitulates knowledge painstakingly accumulated over decades, suggesting that deep learning can discover and formalize biological principles from data.

#### For interpretable machine learning

CytoGem-XAI contributes a case study in designing interpretability into models from the ground up, rather than applying post-hoc explanation methods to black-box models. The three analysis modules—perturbation, hard intervention, and topological weighting—are generalizable beyond metabolism and could be adapted to other graph-structured prediction tasks.

### 4.6 Conclusion

We presented CytoGem-XAI, a hypergraph neural network framework that bridges deep learning and metabolic modeling while providing interpretable, sample-specific analysis. By combining hypergraph representations with perturbation-based carbon source ranking, hard intervention bottleneck identification, and topological pathway attribution, CytoGem-XAI offers biologists actionable insights that directly parallel classical FBA workflows while adding the novel capability of sample-specific characterization. The framework achieves high predictive accuracy (*R*^2^ = 0.862), outperforms existing methods, and recovers known biological principles while generating testable hypotheses. CytoGem-XAI establishes hypergraph neural networks as a versatile paradigm for data-driven metabolic modeling and lays the groundwork for AI-driven discovery in systems biology and metabolic engineering.

## Availability and Implementation

CytoGem-XAI is implemented in Python 3.10 using PyTorch 2.0. The source code, pre-trained models, and all data used in this study are publicly available at: https://github.com/BioCosmos-X/CytoGem-XAI

The repository includes:

- Complete source code with documentation
- Training and evaluation scripts
- Pre-processed datasets (17,400 *E. coli* growth conditions)
- Pre-trained model weights for all 10 cross-validation folds
- Jupyter notebooks for reproducing all figures in this paper
- Usage examples and tutorials
- Docker container for reproducible execution

CytoGem-XAI is released under the MIT license for academic use. Commercial licenses are available upon request.

The iML1515 genome-scale metabolic model of *Escherichia coli* is available from: https://github.com/ecoli/iML1515

The Biolog phenotype microarray data (iML1515_EXP.csv) is included in the repository. All data have been pre-processed as described in Methods 2.1.

For reproducibility, the exact software environment can be reproduced using the provided Dockerfile or the conda environment specification (environment.yml). The code has been tested on Linux (Ubuntu 20.04/22.04) and macOS (12.0+).

## Author Contributions

S.H.C. conceived the study, designed the research, developed the methodology, analysed the data. T.C. analysed the data and wrote the original draft. Z.K.X. collected data and wrote the original draft. L.J.Z., B.G., and J.L.M. administered the project, and supervised the study. All authors reviewed and edited the manuscript and approved the final version.

## Acknowledgements

The authors acknowledge the Biolog database (https://biolog.com) for providing the phenotype microarray data used in this study.

During the preparation of this work, the authors used Kimi (Moonshot AI) and ChatGPT (OpenAI) to assist with language editing and manuscript refinement; Gemini (Google) to generate and refine figures; and ima knowledge base (Tencent) to organize literature and manage references. After using these tools, the authors reviewed, edited, and verified all generated and assisted content as needed and take full responsibility for the accuracy, integrity, and originality of the publication.

## Competing Interests Statement

S.H.C. and T.C. are co-founders of, and hold equity in, BioCosmos X. The other authors declare no competing interests.

## References

[1] Fang, X., Lloyd, C. J. & Palsson, B. O. Reconstructing organisms in silico: genome-scale models and their emerging applications. Nature Reviews Microbiology 18, 731–743 (2020). 10.1038/s41579-020-00440-4.

[2] Henry, C. S. et al. High-throughput generation, optimization and analysis of genome-scale metabolic models. Nature Biotechnology 28, 977–982 (2010). 10.1038/nbt.1672.

[3] Orth, J. D., Thiele, I. & Palsson, B. O. What is flux balance analysis? Nature Biotechnology 28, 245–248 (2010). 10.1038/nbt.1614.

[4] Heirendt, L. et al. Creation and analysis of biochemical constraint-based models using the COBRA Toolbox v.3.0. Nature Protocols 14, 639–702 (2019). 10.1038/s41596-018-0098-2.

[5] Faure, L., Mollet, B., Liebermeister, W. & Faulon, J.-L. A neural-mechanistic hybrid approach improving the predictive power of genome-scale metabolic models. Nature Communications 14, 4669 (2023). 10.1038/s41467-023-40380-0.

[6] Feng, Y., You, H., Zhang, Z., Ji, R. & Gao, Y. Hypergraph Neural Networks. Proceedings of the AAAI Conference on Artificial Intelligence 33, 3558–3565 (2019). 10.1609/aaai.v33i01.33013558.

[7] Yadati, N. et al. HyperGCN: A new method for training graph convolutional networks on hypergraphs. In Advances in Neural Information Processing Systems 32 (2019). NeurIPS proceedings page.

[8] Gao, Y., Zhang, Z., Lin, H., Zhao, X., Du, S. & Zou, C. Hypergraph Learning: Methods and Practices. IEEE Transactions on Pattern Analysis and Machine Intelligence 44, 2548–2566 (2022). 10.1109/TPAMI.2020.3039374.

[9] Chen, C., Liu, Y.-Y. Teasing out missing reactions in genome-scale metabolic networks through hypergraph learning. Nature Communications 14, 2375 (2023). 10.1038/s41467-023-38110-7.

[10] Zhao, Y. et al. A multi-way SMILES-based hypergraph inference network for metabolic model reconstruction. Communications Biology 9, 531 (2026). 10.1038/s42003-026-09761-1.

[11] Gao, Y., Feng, Y., Ji, S. & Ji, R. HGNN+: General Hypergraph Neural Networks. IEEE Transactions on Pattern Analysis and Machine Intelligence 45, 3181–3199 (2023). 10.1109/TPAMI.2022.3182052.

[12] Chien, E., Pan, C., Peng, J. and Milenkovic, O. You are AllSet: A Multiset Function Framework for Hypergraph Neural Networks. In International Conference on Learning Representations (2022). https://openreview.net/forum?id=hpBTIv2uy_E.

[13] Monk, J. M. et al. iML1515, a knowledgebase that computes Escherichia coli traits. Nature Biotechnology 35, 904–908 (2017). 10.1038/nbt.3956.

[14] Gilmer, J., Schoenholz, S. S., Riley, P., Vinyals, O. & Dahl, G. E. Neural Message Passing for Quantum Chemistry. In Proceedings of the 34th International Conference on Machine Learning, 1263–1272 (2017). https://proceedings.mlr.press/v70/gilmer17a.html.

[15] Xiong, R. et al. On Layer Normalization in the Transformer Architecture. In Proceedings of the 37th International Conference on Machine Learning, 10524-10533 (2020). https://proceedings.mlr.press/v119/xiong20b.html.

[16] Velickovic, P. et al. Graph Attention Networks. In International Conference on Learning Representations (2018). https://openreview.net/forum?id=rJXMpikCZ.

[17] Srivastava, N., Hinton, G., Krizhevsky, A., Sutskever, I. & Salakhutdinov, R. Dropout: A simple way to prevent neural networks from overfitting. Journal of Machine Learning Research 15, 1929–1958 (2014). https://www.jmlr.org/papers/v15/srivastava14a.html.

[18] Loshchilov, I. & Hutter, F. Decoupled Weight Decay Regularization. In International Conference on Learning Representations (2019). https://openreview.net/forum?id=Bkg6RiCqY7.

[19] Paszke, A. et al. PyTorch: An imperative style, high-performance deep learning library. In Advances in Neural Information Processing Systems 32, 8024–8035 (2019). NeurIPS proceedings page.

[20] Chen, T. & Guestrin, C. XGBoost: A scalable tree boosting system. In Proceedings of the 22nd ACM SIGKDD International Conference on Knowledge Discovery and Data Mining, 785–794 (2016). 10.1145/2939672.2939785.

